# Enhancing burst activation and propagation in cultured neuronal networks by photo-stimulation

**DOI:** 10.1101/027177

**Authors:** Ghazaleh Afshar, Ahmed El Hady, Theo Geisel, Walter Stuehmer, Fred Wolf

## Abstract

Spontaneous bursting activity in cultured neuronal networks is initiated by leader neurons, which constitute a small subset of first-to-fire neurons forming a sub-network that recruits follower neurons into the burst. While the existence and stability of leader neurons is well established, the influence of stimulation on the leader-follower dynamics is not sufficiently understood. By combining multi-electrode array recordings with whole field optical stimulation of cultured Channelrhodopsin-2 transduced hippocampal neurons, we show that fade-in photo-stimulation induces a significant shortening of intra-burst firing rate peak delay of follower electrodes after offset of the stimulation compared to unperturbed spontaneous activity. Our study shows that optogenetic stimulation can be used to change the dynamical fine structure of self-organized network bursts.

## Introduction

Synchronized bursting is a major constituent of spontaneous activity that has been observed in cultured hippocampal and cortical neurons [1, 2]. There are numerous quantitative studies that investigated the ignition and spread of collective spontaneous bursting activity [3,4,5,6,7] showing that the order of activation within a synchronized burst is a stereotypical hierarchical process [4] and that multiple ignition sites, termed as initiation zones [6,7], privileged neurons [4]or leader neurons [5], create network bursts by recruiting follower neurons. Leader neurons are relatively robust and they carry information about the identity of the burst. They are supposed to be part of an underlying sub-network that is excited first [5], which then recruits the follower neurons into the orchestrated activation of neuronal cell assemblies. Functionally, leader-follower neuron temporal relationships reflect the dynamical state of the network and can be used to assess network topology [3]. Despite numerous reports on the existence of leader and follower neurons, the effect of stimulation modifying the leader – follower dynamics have not been sufficiently studied. The modification of leader – follower relationships may provide insights on how the internal structure of synchronized bursting activity can be regulated by activity dependent network-level potentiation. This will contribute to our understanding of the relationship between network level plasticity and neuronal recruitment into synchronized bursting activity. Therefore, we investigated the leader-follower dynamics using a combination of multielectrode array recording and optical stimulation of channelrhodopsin-2 transduced hippocampal cultures. This approach offers a non-invasive technique for recording and stimulating cultured neuronal networks. Electrodes are divided into two subsets of 1) leader electrodes, which are a small subset of neurons initiating the bursts and 2) follower electrodes, which register other neurons being recruited by the leaders into an emerging synchronized activity burst. We found that a fade-in photo-stimulation paradigm consisting of slowly rising ramps of blue light induced a shortening of the intra-burst firing rate peak delay after stimulation reflecting a tightening of leader-follower temporal relationships. Interestingly, the leader– follower dynamics are differentially modulated by different photo-stimulation paradigms as we found that fade-in stimulation can substantially affect leader-follower dynamics more potentially than pulsed stimulation. These Results raise the possibility to modulate the intrinsic dynamics of self-organized bursting activity and to study how activity dependent plasticity can modify it.

### Experimental details

#### Cell culture, transduction and multi-electrode array recordings

Hippocampal neurons isolated from E18 Wistar U rats were cultured following primary hippocampal culture procedures described in [8] and plated on multi-electrode arrays (MEA; type TiN-200-30iR from Multichannel Systems, Reutlingen, Germany) at a density of 1000 cells per mm^2^. The multi-electrode arrays were coated with 1ml of a mixture, composed of 600 μl poly-D-lysine (50μg / ml) and 200 μl (10μg / ml) laminin dissolved in 15 ml bidistilled water, before plating the cells on it. All animals were kept and bred in the animal house of the Max Planck Institute for Experimental Medicine according to the German guidelines for experimental animals. All experimental procedures were carried out with authorization of the responsible federal state authority. The MEAs were covered with the Teflon fluorinated ALA-science caps (ALA scientific instruments, US). The cells were kept in an incubator at 37°C, 8% CO2 and 90 % humidity. 14 days in vitro cultures were transduced with AAV-CAG-ChR2-YFP virus [9, 10]. Recordings were done after 21 days in vitro. The recordings were made on a 60 channel MEA amplifier (MEA-1060 Inv, Multichannel Systems, Reutlingen, Germany). Data from MEAs were captured at 25 kHz using a 64-channel A/D converter and MC_Rack software (Multichannel Systems, Reutlingen, Germany). After high pass filtering (Butterworth 2nd order, 100 Hz) extracellular action potential waveforms were detected in a cutout recorded 1 ms before and 2 ms after crossing a threshold of -20 μV, which was > 3 times standard deviations of the baseline activity.

#### Whole field blue light stimulation

Two protocols of whole field blue light stimulation were used: (1) 40 repetitions of 1 second rectangular (pulsed) light pulses and (2) fade-in stimulation designed as 40 repetitions of slowly ramped light waveform up to the level of constant pulses with frequency of 0.5 Hz. With both pulsed and fade-in stimulation, twelve experiments on twelve cultures were performed. In each experiment, before the onset of the stimulation, the spontaneous activity of the culture was recorded for 5 minutes. Then the culture was stimulated with one of the two stimulation protocols. After offset of stimulation spontaneous activity was recorded for 12 minutes.

#### Network Dynamics Data analysis

Quantification of burst dynamics was restricted to the subset of active electrodes. *Active electrodes* (AE) were defined as electrodes that had a spontaneous firing rate of more than 0.1 Hz.

#### Burst detection

For burst detection we modified the method suggested by [11]. Bursts were defined as sequences of at least two spikes with all inter-spike intervals lower than a threshold value. The threshold was defined as 1/4 of the inverse average firing rate of all AEs. After detecting bursts on all AEs, they were sorted in temporal order. A synchronized burst was defined as a group of bursts across several electrodes that overlapped in time. After detecting all synchronized bursts, the synchronized bursts that were separated by less than 5/4 of the threshold, inter-spike intervals were merged into one synchronized burst.

#### Peri-stimulus time histograms (PSTHs)

were calculated using a 20 msec time bin. The level of activity of individual cultures was characterized by the corresponding spontaneous average firing rate, which varied from culture to culture. The average PSTH was obtained from the PSTHs of each experiment normalized with the spontaneous average firing rate before stimulus of the corresponding culture. The time course of the average firing rate during pulsed stimulation was markedly different from that induced by fade-in stimulation. This difference can be seen in the averaged normalized peri-stimulus time histogram (PSTH) plots shown in Fig. 1d.

**Fig. 1:**
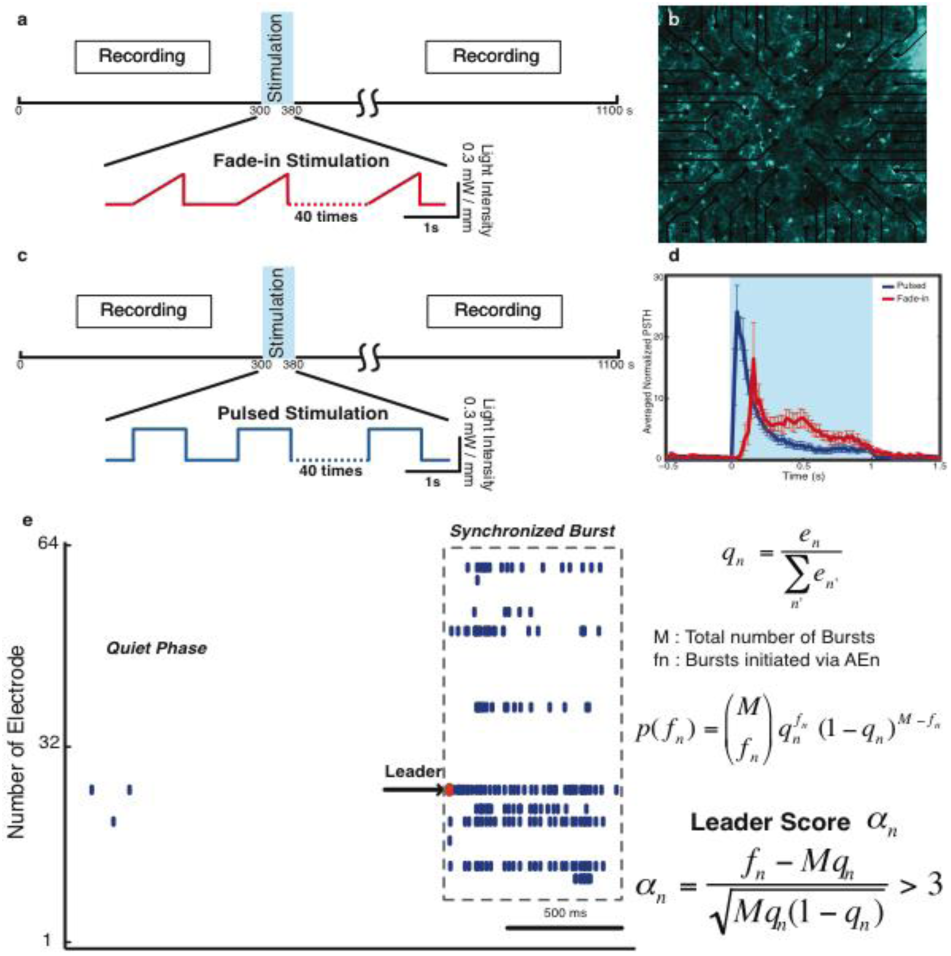
Experimental setting. Panel (**a**) depicts the recording and photo-stimulation design with fade-in stimulation. Panel (**b**) shows the Channelrhodopsin-2 transduced neurons cultured on multi-electrode array. Panel (**c**) depicts the recording and photo-stimulation design with pulsed stimulation. Panel presents the electrode averaged normalized peri-stimulus time histogram (PSTH) for both pulsed (dark blue) and fadein stimulation (red). In panels (**e**) the leader detection method is illustrated by using an example of one burst. The first spike fired from the leader electrode is marked in red.

#### Intra-burst firing rate (IBFR)

In order to obtain the IBFR, the spike trains of AEs for each synchronized burst were convolved with a Gaussian kernel of standard deviation of 5 ms and averaged over all bursts at each AE.

#### Leaders and followers

To qualify the leader electrodes, we modified the method suggested by [5]. This modification is done due to the different burst detection method used in their study. In study [5], synchronized bursts were divided into two classes of bursts with or without pre-burst and for leader detection, only bursts with pre-bursts are considered. However, in our study we consider all detected synchronized burst.

Let M be the total number of detected synchronized bursts. For each AE_n_, *e*_*n*_ is defined as the total number of evoked spikes in inter-burst intervals,

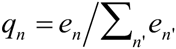

is the relative firing rate in quiet phase and *f*_*n*_ is the actual number of bursts initiated via AE_n_. The probability of leading the fraction of *f*_*n*_ out of M bursts would be given by the following binomial distribution:

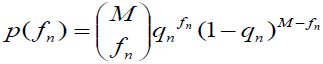

This equation gives the probability of firing first in the burst if firing order was completely random. Statistically this null hypothesis can be rejected if *f*_*n*_ is three times standard deviations above the natural expectation value. Therefore, an AE_n_ is defined as a leader if the leadership score *αn* fulfills the following condition:

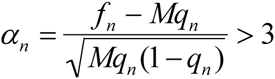

If a leader electrode has led less than 3% of the bursts, it’s excluded from being a leader. Besides the detected leader electrodes before and after stimulus, the rest of the electrodes were denoted as followers.

#### Peak delay

was defined as the time delay of the first peak which larger than 2/3 of *maximum peak of IBFR* at each electrode from onset of the burst.

## Results

Our experimental setup combines multichannel recording using multi-electrode arrays and whole field photo-stimulation. Fig. 1b shows a 21 DIV Channelrhodopsin-2 transduced embryonic hippocampal neurons plated on 60 channels multi-electrode array (MEA). The experimental paradigm and the used photo-stimulation protocols (1) fade-in and (2) pulsed are presented in Fig. 1a,c. During the stimulation, the time course of the average firing rate during pulsed stimulation was different from that induced by fade-in stimulation. The difference can be seen in the average normalized peri-stimulus time histogram (plot shown in Fig. 1d).

The leader detection method is illustrated in Fig. 1e. Performing leader analysis on our dataset, we could confirm that leader electrodes exist in our recordings and can be detected. The alpha scores of all AEs before and after fade-in and pulsed stimulation are shown in Fig. 2a,b. As it is shown in Fig. 2a,b, the leader electrodes were mainly robust before and after stimulation. Most of the bursts were initiated via one of the leader electrodes. On average (mean±SD) 68% ± 21 and 63% ± 18 of all bursts were initiated via leader electrodes before and after fade-in stimulation. In case of pulsed stimulation, 70% ± 15 and 70% ± 15 of all bursts were initiated via leader electrodes before and after stimulation. Averaging over all experiments (n = 12), we found that 2.2% ± 1.9 electrodes were leaders before and after fade-in stimulation and 1.9% ± 0.8 electrodes were leaders before and after pulsed stimulation (Fig. 2c). In order to study the effect of stimulation on leader and follower electrodes, only the bursts initiated via one of the leader electrodes were taken into account for the whole analysis.

**Fig. 2:**
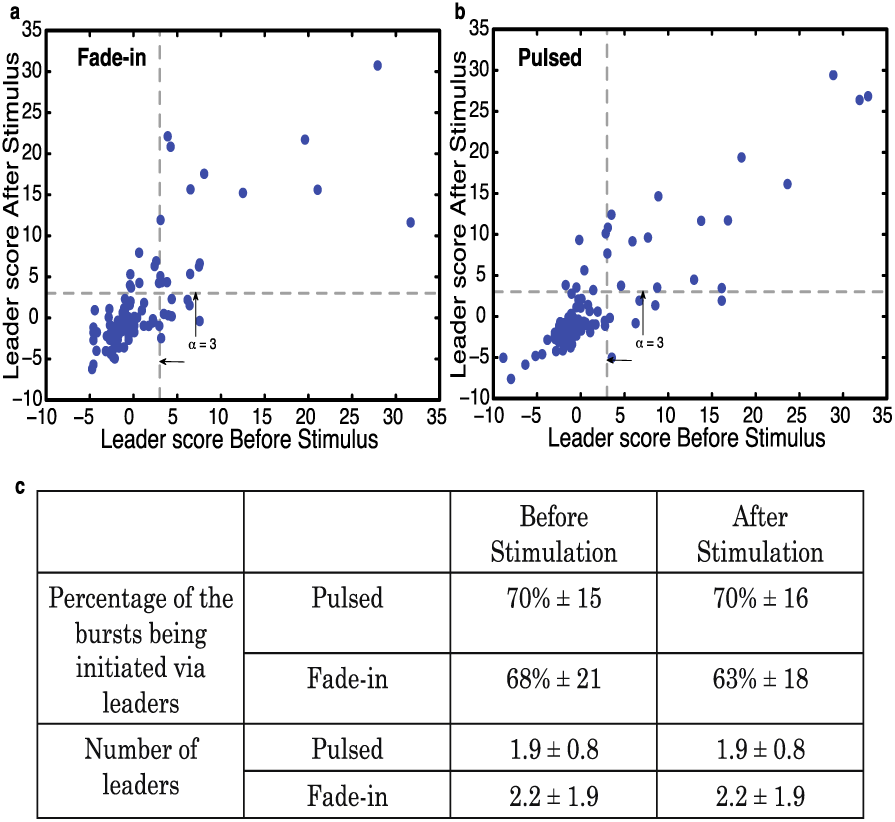
Leader-follower statistics. Panels (**a,b**) show the alpha-score scatter plot after versus before fade-in and pulsed stimulation. Panel (**c**) summarizes the statistics of leader electrodes before and after fade-in and pulsed stimulation

We investigated the intricate details of the network level enhanced activity by comparing the maximum peak of the IBFR at each electrode before and after stimulation. This analysis was performed for both leader and follower electrodes. In case of fade-in stimulation (Fig. 3c), the maximum Afshar *et al*. peak of IBFR after stimulation for 125 follower and 35 leader electrodes increased significantly compared to unperturbed spontaneous activity (followers and leaders respectively: p=0.01, p<10^-2^). As for pulsed stimulation (Fig. 3e), the maximum peak of IBFR after stimulation for 132 follower and 29 leader electrodes also increased substantially compared to unperturbed spontaneous activity prior to stimulation (followers and leaders respectively: p<10^-5^, p=0.02). Moreover, cumulative distribution of the ratio of the maximum peak of IBFR after to before stimulation showed a significant difference b etween fade-in and pulsed stimulatio. 3g) (p=0.02).

**Fig. 3:**
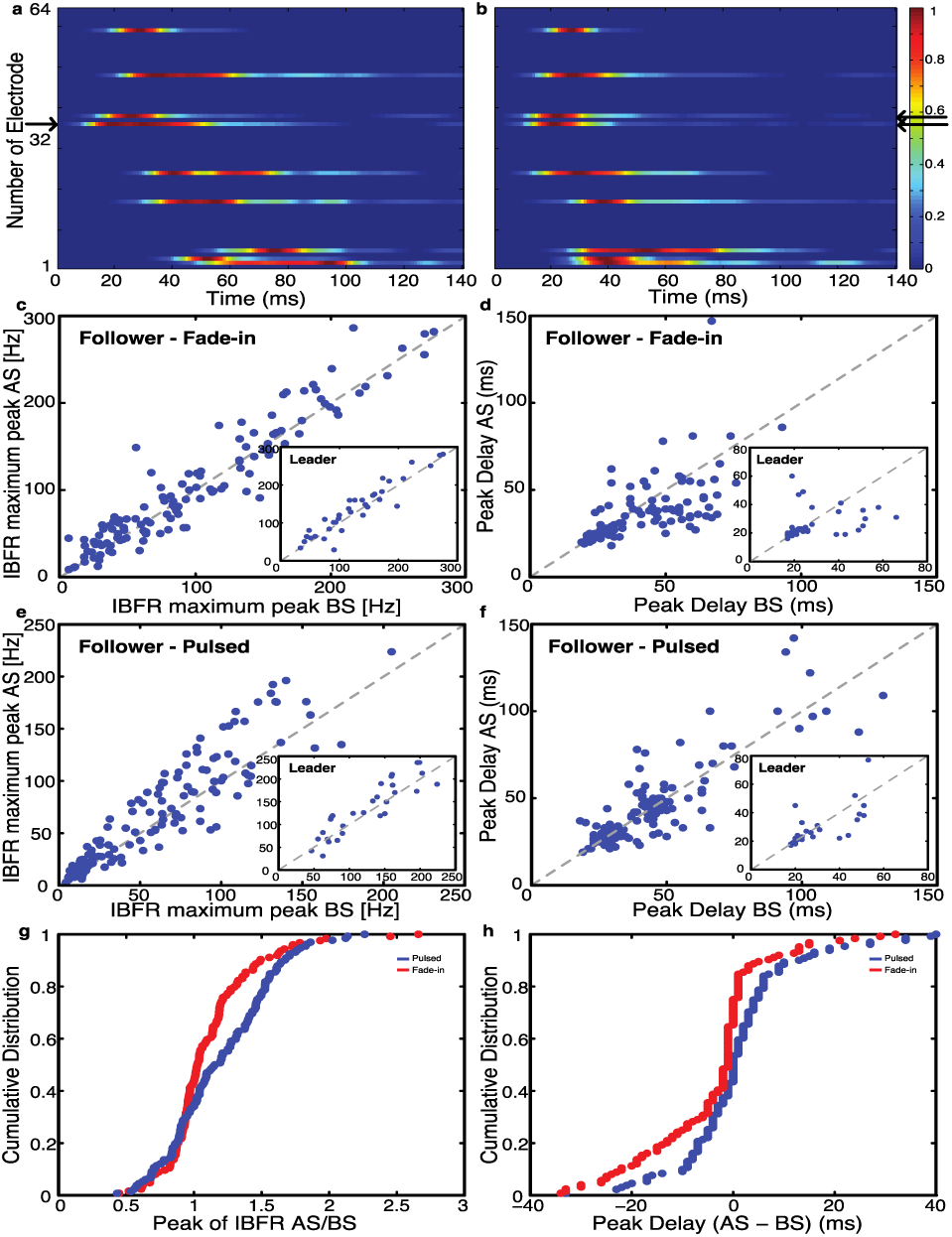
Leader-follower dynamics. The average normalized IBFR over all bursts of one experiment before and after fadein stimulation are shown in panels (**a,b**). The leader electrode is marked by a black arrow. **Panel (c)** shows IBFR maximum peak of follower and leader electrodes (inset) after fade-in stimulus compared to before stimulation. IBFR maximum peak increase significantly after fade-in stimulation for followers and leaders respectively (p=0.01, p<10^-2^). **Panel (d)** shows the IBFR peak delay after fade-in stimulus compared to before stimulation. The IBFR peak delay decreases significantly compared to before stimulus (p<10^-5^) and no significant difference in case of leader electrodes (inset) (p>0.05). **Panel** shows IBFR maximum peak of follower and leader electrodes (inset) after pulsed stimulation compared to before stimulation. IBFR maximum peak increase significantly after pulsed stimulation for followers and leaders respectively (p<10^-5^, p=0.02). **Panel (f)** shows the IBFR peak delay after pulsed stimulation compared to before stimulation. The IBFR peak delay has no significant change either for followers or leaders (inset) (p>0.05). **Panel (g)** shows the cumulative distribution of the ratio of the maximum peak of IBFR after to before stimulation for both fade-in stimulus (red line) and pulsed stimulus (blue line). There is a significant difference between fade-in and pulsed stimulation cumulative distribution of the ratio of the maximum peak of IBFR after to before stimulation (p=0.02). **Panel (h)** shows the cumulative distribution of the difference of the IBFR peak delay after to before stimulation for both fade-in stimulus (red line) and pulsed stimulus (blue line). There isa significant difference between fade-in and pulsed peak delay difference after and before (p<10^-2^). For pair comparisons the Wilcoxon signed rank test and in case of cumulative distributions the Wilcoxon rank sum test is used.

We next compared the time delay of the IBFR peaks from the onset of the burst. The normalized IBFRs of AEs over all bursts initiated via the leader electrodes of one experiment before and after fade-in stimulation shows a shorter peak delay after stimulation (Fig. 3a,b). In Fig. 3d, the peak delay scatter plot of 125 follower electrodes from 12 experiments shows that the peak delay of follower electrodes gets significantly shorter after fade-in stimulation compared to the unperturbed spontaneous activity (p<10^-5^). However, in case of leader electrodes (35 electrodes in total from 12 experiments) no significant change in peak delay after stimulation is observed (Fig. 3d Inset). In case of pulsed stimulation, no significant change in peak delay of follower and leader electrodes was observed after offset of the stimulation compared to unperturbed spontaneous activity (Fig. 3f) (132 follower electrodes and 29 leader electrodes from n=12 experiments) (p>0.05 in both cases). In order to compare the change of peak delays after offset of the stimulation between pulsed and fade-in stimulation, the cumulative distribution of the difference between peak delay after and before stimulation was used (Fig. 3h). This shows that, with fade-in stimulation the decrease in peak delays of IBFRs after stimulation is much more pronounced compared to pulsed stimulation (p<10^-2^).

## Discussion

Using an experimental setup combining multielectrode array recording with optogenetic stimulation designed as either fade-in or pulsed photo-stimulation, we were able to modify the propagation of activity at burst onset across neurons. As the collective network dynamics is dominated by bursts, the intricate structure of the bursts likely reflect directed interactions between neurons and therefore are expected to change as a result of stimulation. In this study, we found that the leader–follower electrode temporal relationship gets tightened after fade-in photo-stimulation. We found that leader – follower dynamics is more profoundly modulated by fade-in stimulation than by pulsed stimulation.

Synchronized burst initiation involves two distinct processes: the activation of first-to-fire neurons and the recruitment of follower neurons by “leader neurons” into the burst. The activation process of the bursts has been reported to be a stereotypical process involving “leader neurons” that recruit “follower neurons” to participate in the burst[4]. In our recordings, “leader neurons” were mainly robust, which has been confirmed in other studies that pinpointed to the robustness of the first-to-fire neurons. Moreover, most of the bursts originated via one of the leader electrodes (∼70% both before and after stimulation), which is also in agreement with previous reports [5].The changes that were induced in the network, upon photostimulation, are reflected in the relationship between leader and follower neurons and not the identity of leader and follower neurons per se. Leader neurons might alternatively be regarded as a kind of hub neurons. Previous reports described that hub circuits can be involved in the initiation of population bursts in cortical slice cultures [12]. Moreover, hub neurons have been reported to form a connected network that initiate synchronized bursting [13] in the same manner in which leader neurons have been proposed to form a distinct sub-network among themselves [5].

We have found that the normalized IBFRs of AEs Afshar *et al*. over all bursts initiated via the leader electrodes of the same experiment before and after fade-in stimulation shows a shorter peak delay after stimulation reflecting that the follower neurons fire much faster in relation to the leader neurons, thus they have a tighter relationship with the leader in the neuronal recruitment sequence. In case of pulsed stimulation, no significant change in peak delay of follower and leader electrodes is observed after offset of the stimulation compared to unperturbed spontaneous activity reflecting the fact that the temporal relationship between leader and follower neurons within the recruitment sequence is not affected. The aforementioned indicates that the follower – leader relationship is differentially regulated by photostimulation paradigm design. The tightening of the leader–follower relationship observed with fade-in photostimulation might result from synaptic strengthening following a spike-timing dependent plasticity mechanism [14-16] owing to the enhancement of inter-neuronal correlations resulting from fade-in stimulation. It might also result from the increased number of spikes within the bursts as a result of photo-stimulation leading to the potentiation of network intrinsic dynamics as evidenced by the significant increase in the maximum peak of the average intra-burst firing rate after stimulation. In the case of pulsed stimulation, the stimulation might essentially override the network activity thus preventing the enhancement of the networks’ intrinsic dynamics. Our results thus highlight that stimulation paradigms need to be tailored to respect and activate an neuronal circuits intrinsic dynamics in order to effectively engage plasticity mechanisms.

***

## Acknowledgement

The authors would like to thank H. Kerren, A. Wallach, U. Egert, M. Giugliano, M. Gutnick, E. Moses, A. Neef, G. Rapp, S. Solla, and S. Shoham for discussions. Ahmed El Hady was supported by a Neurosenses doctoral fellowship and Ghazaleh Afshar by an International Max Planck Research School doctoral fellowship. The authors acknowledge the BMBF for funding (Grant number 01GQ0811, 01GQ01005B, 01GQ0922), ZIM grant (KF2710201 DF0), the DFG (SFB 889 and Cluster of Excellence “Nanoscale Microscopy and Molecular Physiology of the Brain”), and VolkswagenStiftug (ZN2632). The European Neuroscience Institute-Göttingen is jointly funded by the Max Planck Society and University Medicine Göttingen.

